# Achronopresence: how temporal visuotactile and visuomotor mismatches modulate embodiment

**DOI:** 10.1101/596858

**Authors:** Marte Roel Lesur, Marieke Lieve Weijs, Colin Simon, Oliver Alan Kannape, Bigna Lenggenhager

**Author notes:** Corresponding authors: Marte Roel Lesur, Binzmühlestrasse 14, Box 9, Zurich, 8050, Switzerland. and Bigna Lenggenhager, Binzmühlestrasse 14, Box 9, Zurich, 8050, Switzerland. = Shared first authorship (Marte Roel Lesur, Marieke Lieve Weijs).

## Abstract

The loss of body ownership, the feeling that your body and its limbs no longer belong to you, presents a severe clinical condition that has proven difficult to study directly. We here propose a novel paradigm using mixed reality to interfere with natural embodiment using temporally conflicting sensory signals from the own hand. In Experiment 1 we investigated how such a mismatch affects phenomenological and physiological aspects of embodiment, and identified its most important dimensions using a principle component analysis. The results suggest that such a mismatch induces a strong reduction in embodiment accompanied by an increase in feelings of disownership and deafference, which was, however, not reflected in physiological changes. In Experiment 2 we refined the paradigm to measure perceptual thresholds for temporal mismatches and compared how different multimodal, mismatching information alters the sense of embodiment. The results showed that while visual delay decreased embodiment both while actively moving and during passive touch, the effect was stronger for the former. Our results extend previous findings as they demonstrate that a sense of disembodiment can be induced through controlled multimodal mismatches about one’s own body and more so during active movement as compared to passive touch. Based on the ecologically more valid protocol we propose here, we argue that such a sense of disembodiment may fundamentally differ from disownership sensations as discussed in the rubber hand illusion literature, and emphasize its clinical relevance. This might importantly advance the current debate on the relative contribution of different modalities to our sense of body and its plasticity.

## 1. Introduction

Over the past two decades, experimental evidence has shown that the sense of body of healthy subjects is remarkably plastic and built upon a constant prediction, weighting and integration of multimodal signals (e.g. Blanke, 2012). Protocols involving multimodal stimulation suggest that a majority of healthy individuals embody foreign or virtual limbs or full bodies when bodily sensations (e.g. body movements or touch) are visually displayed in synchrony to matching sensations on the hidden body (e.g. Botvinick & Cohen, 1998; Tsakiris, Prabhu, & Haggard, 2006; Lenggenhager, Tadi, Metzinger, & Blanke, 2007; Slater, Spanlang, Sanchez-Vives, & Blanke, 2010). Such illusory embodiment is usually manifested by the senses of body ownership and agency (Kalckert & Ehrsson, 2012; Tsakiris et al., 2006), and has been evidenced using a variety of experimental setups using both explicit (i.e. questionnaires) and implicit (i.e. proprioceptive drift or physiological responses) measures (Blanke, Slater, & Serino, 2015).

This line of research predominately investigated the influence of multimodal coherence on illusory embodiment of an external or supernumerary bodily object; far more elusive, however, is how breaking multimodal information about the own body might reduce embodiment or even induce a feeling of disembodiment (Gentile, Guterstam, Brozzoli, & Ehrsson, 2013; Graham, Martin-Iverson, Holmes, & Waters, 2015; Hoover & Harris, 2012; Kannape, Smith, Moseley, Roy, & Lenggenhager, 2019; Newport & Preston, 2011; Otsuru et al., 2014). This is surprising as disorders of bodily self awareness in clinical populations predominantly manifest in a loss of embodiment, as a break of (own) body ownership and one’s sense of agency (Aglioti, Smania, Manfredi, & Berlucchi, 1996; Brugger & Lenggenhager, 2014; Otsuru et al., 2014; Vallar & Ronchi, 2009). For example, in the case of somatoparaphrenia, resulting from a brain lesion, patients lack the feeling of ownership for the contralesional arm, often attributing that arm to someone else (Aglioti et al., 1996; Brugger & Lenggenhager, 2014; Vallar & Ronchi, 2009) or even showing aggression towards it (Lötscher, Regard, & Brugger, 2006). Similarly, individuals suffering from body integrity dysphoria feel strong alienation from one or several body parts often combined with a desire for amputation (Blom, Hennekam, & Denys, 2012; Brugger & Lenggenhager, 2014; Lenggenhager, Hilti, & Brugger, 2015). Such a feeling of disembodiment might also extend to the full body, both in neurological (Smit, van Stralen, van den Munckhof, Snijders, & Dijkerman, 2018) as well as in psychiatric disorders, like during depersonalization (Davidson, 1966; Sierra, Baker, Medford, & David, 2005).

Important theoretical differences between ownership of an external body, reduced ownership for one’s own body, and body *dis*ownership have been proposed (de Vignemont, 2011), and the degree of alteration in embodiment of the own body in illusory limb of full body ownership paradigms remains elusive. While some authors suggest decreased ownership for the real body based on questionnaire (Longo, Schüür, Kammers, Tsakiris, & Haggard, 2008; Moseley et al., 2008) or even immunological data (Barnsley et al., 2011), others found disownership of one’s own body to be rare and rather weak in rubber hand illusion like setups (Folegatti, Vignemont, Pavani, Rossetti, & Farnè, 2009). Data from individuals with clinically caused alterations leading to loss of own-body ownership generally suggest enhanced illusory ownership for an external body, pointing to different mechanisms between embodiment and disembodiment in patients suffering from schizophrenia (Thakkar, Nichols, McIntosh, & Park, 2011; see Shaqiri et al., 2018 for alternative findings during full body illusions), body integrity dysphoria (Lenggenhager et al., 2015) or somatoparaphrenia (Smit et al., 2018; van Stralen, van Zandvoort, Kappelle, & Dijkerman, 2013; White & Aimola Davies, 2017). This is further evidenced by a voxel-based lesion symptom mapping study that found a partial dissociation between brain areas involved in own-limb disembodiment as compared to supernumerary embodiment (Martinaud, Besharati, Jenkinson, & Fotopoulou, 2017).

Here we directly manipulated embodiment of one’s biological hand using a controlled multisensory conflict, without the use of a proxy/rubber hand. Previous studies suggest a feeling of disownership and numbness during delayed and therefore conflicting visual feedback of a tactile event in a mixed reality setup using infrared camera feed (Kannape et al., 2019) or prerecorded video (Gentile et al., 2013). By using an online, naturally colored, and large field of view video-feed on a head mounted display (HMD), which provides a direct view on the own body in its current environment, we adapted such setups to be more realistic and thus more ecologically valid. Our setup was created to induce a strong prior assumption of actually viewing one’s own body. We then manipulated the delay of the video feed digitally, thus controlling the latency of visual as compared to other bodily signals (i.e. visuomotor and visuotactile or potentially others). We used this setup in two different experiments to evaluate the relative influence of multimodal mismatch about one’s own body on the sense of embodiment and its physiological correlates. Importantly, while previous studies investigated visuotactile incongruency, they did not account for differential sensitivity of various multimodal coherences in the construction and maintenance of the bodily self. Yet, differential roles of motor and somatosensory signals in the sense of body have been suggested (Asai, 2015; Tsakiris, Longo, & Haggard, 2010; Tsakiris et al., 2006) and the role of actively (moving) in comparison to passively perceiving bodily signals to the bodily self has been extensively discussed (Grechuta, Ulysse, Rubio Ballester, & Verschure, 2019; Pia, Garbarini, Kalckert, & Wong, 2019).

In Experiment 1 we manipulated visuotactile coherence, which is classically used to induce altered embodiment in rubber hand illusion-like paradigms. For two stimulation durations (1 and 3 minutes) the participant’s hand was stroked with a paintbrush while the visual feedback, was either delayed (∼illusion condition) or not (∼control condition). Alterations in embodiment, ownership, sensations of deafferentation and related phenomenological sensations were measured using questionnaires adapted from (Botvinick & Cohen, 1998; Kannape et al., 2019; Lenggenhager et al., 2007; Longo et al., 2008). Furthermore, previously suggested implicit correlates of embodiment, namely skin temperature (see Moseley et al., 2008, but see also de Haan et al., 2017 for a critical view) and skin conductance responses (SCR) to threat (see Armel & Ramachandran, 2003) were assessed. Heart rate variability (HRV) measures were added, as homeostatic processes have suggested to be altered in conditions of alteration in body ownership (Barnsley et al., 2011). A measure of interoceptive accuracy has been included as poor accuracy has previously shown to be related to higher susceptibility to illusory ownership and thus a more plastic bodily self (Monti, Porciello, Tieri, & Aglioti, 2019; Tsakiris, Jiménez, & Costantini, 2011). We hypothesized that the sensory conflict between tactile and delayed visual feedback would result in a reduced sense of embodiment and enhanced sense of disembodiment, which would be reflected in both explicit (subjective) and implicit (physiological) measures, especially in participants with a weak interoceptive accuracy.

In Experiment 2, we investigated the temporal thresholds for detecting synchrony for visuomotor as compared to visuotactile delays and how different delays relate to the feeling of *dis*embodiment. While, systematic empirical comparisons remain rather scarce, some studies have suggested the relative importance of active movements versus passive touch for an integrated and global sense of body (Burin et al., 2015; Tsakiris et al., 2010, 2006); and there is some evidence that sensory and motor signals may contribute differently to illusory embodiment with visuomotor synchrony being more important for illusory embodiment than visuotactile synchrony (Kokkinara & Slater, 2014; Roel Lesur, Gaebler, Bertrand, & Lenggenhager, 2018). Moreover, there is evidence suggesting that multisensory integration of peripheral signals behaves differently when followed by efferent signals as compared to afferent signals (Zierul, Tong, Bruns, & Röder, 2018).

## 2. Experiment 1

### 2.1 Method

#### 2.1.1 Participants

Thirty healthy volunteers participated in Experiment 1 (10 males; 25 ± 3.8 years). Participants provided informed consent and received either course credit or financial compensation.

For the principal component analysis (PCA) of the questionnaire responses after synchronous versus asynchronous stroking, we additionally included the participants of Experiment 2 (see 3.1.1), as well as 15 participants (3 males; 22.2 ± 2.4 years) from a previously unpublished experiment resulting in a total of 77 participants (27 males; 22.9 ± 4.0 years).

All protocols were approved by the Ethics Committee of the Faculty of Arts and Social Sciences at the University of Zurich (Approval Number 17.12.15). The studies were performed in accordance with the ethical standards of the Declaration of Helsinki.

#### 2.1.2 Apparatus for stimulation

An Oculus CV1 HMD (Oculus VR, Irvine, CA, USA) was used for the visual stimulation. An ELP 180° webcam (Ailipu Technology Co., Ltd, Guangdong, China) was positioned on the front of the HMD, set to 30 frames per second and resolution of 1024 × 768 pixels. The camera was rotated 90° to have the wide field of view on the vertical axis in order to show the full-body. The control system was designed using Unity 2017 for delaying the camera feed, rotating the image, mapping it to a 3D model approximately matching the distortion of the camera-lens, and projecting the image on the HMD. The questionnaires and randomization were also built within Unity 2017. The system was run on an Alienware 15 R3 computer (Nvidia Geforce GTX 1080 8GB; 16GB RAM; Intel Core i7; Windows 10), which added a mean intrinsic delay of 139.1ms (*SD* = 18.3 ms).

#### 2.1.3 Procedure

##### 2.1.3.1 Heartbeat counting task

At the beginning of the experiment, participants performed a heartbeat counting task (Schandry, 1981), see Figure 1 for general procedure and order of the experiment. Participants were instructed to count their heartbeats, without taking their pulse. They were informed that the time of the intervals would vary, to prevent them relying on time estimation instead of actual counting of the heartbeats. Three intervals of 25, 35, and 45 s were presented in randomized order, and the start and end of each interval was indicated by a tone. During the task, electrocardiograms (ECG) were recorded with a Biopac MP150 system and ECG100C amplifier (Goleta, USA) at 1000 Hz sampling rate. Three ECG electrodes (Red Dot, 3M, Neuss, Germany) were placed on the left and right clavicle and on the lowest left rib. The electrodes were left to measure ECG throughout the experimental procedure.

**Figure 1:**
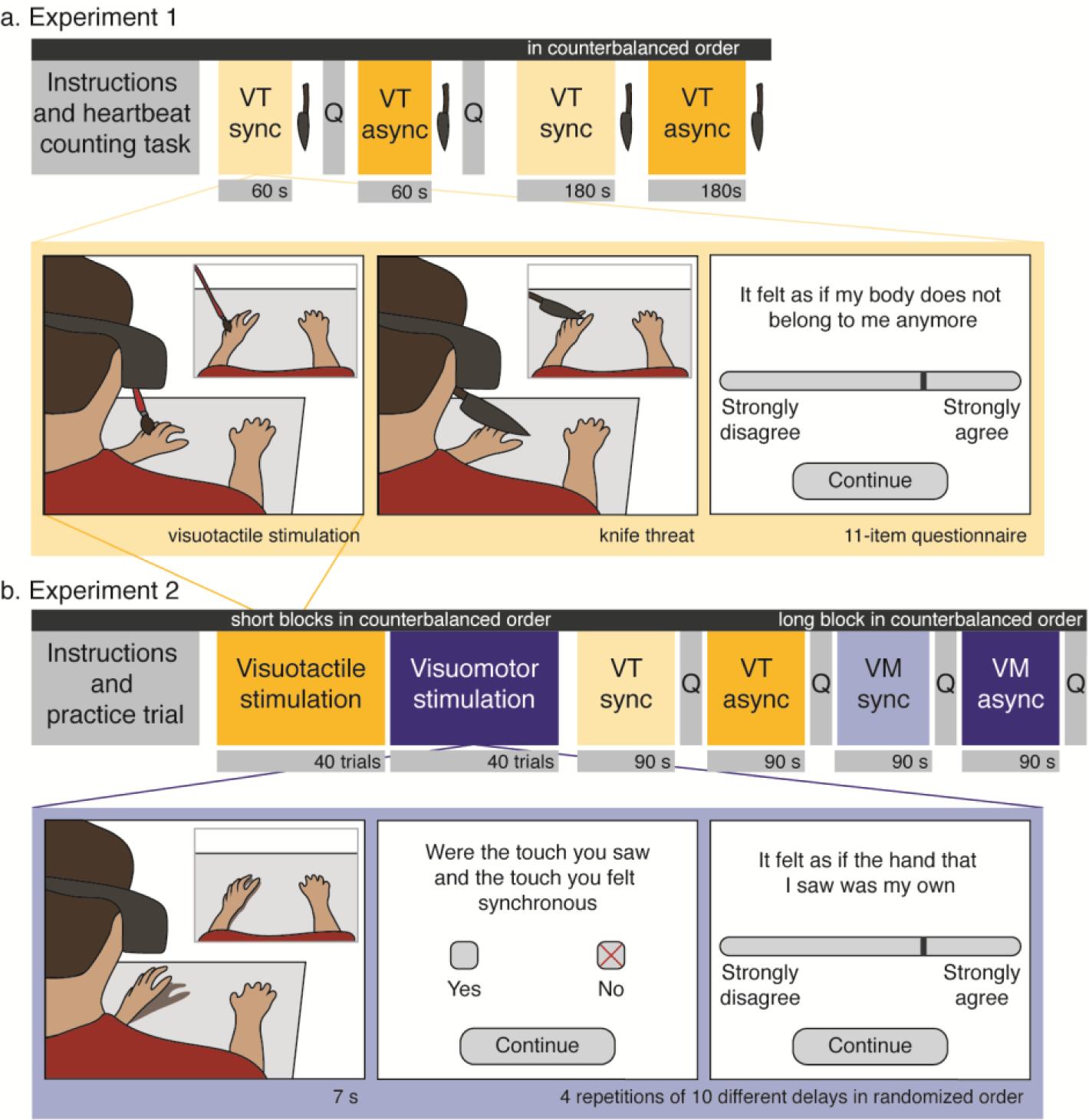
Experimental setup in (a) Experiment 1 and (b) Experiment 2. In Experiment 1, the visuotactile stimulation was either synchronous (VT sync) or asynchronous (VT async). Each stimulation was followed by a knife threat, and in the 60 s blocks also by the embodiment questionnaire (Q). In Experiment 2 the visuotactile stimulation was similar to that of Experiment 1, but this time, as with the visuomotor stimulation, was presented for 7 s. After each trial two questions appeared on the HMD. This was repeated 40 times in each modality, with four repetitions of 10 possible delay steps. Then, a long block followed with synchronous visuotactile (VT sync) and visuomotor (VM sync) as well as asynchronous visuotactile (VT async) and visuomotor (VM async) stimulation.

The heartbeat perception score was calculated as 1/3 Σ (1-|recorded heartbeats – perceived heartbeats| / recorded heartbeats), so that higher scores indicate higher accuracy. Data from 10 participants were excluded due to technical difficulties with the ECG recording equipment, missing markers, or because they did not understand the task.

##### 2.1.3.2 Visuotactile stimulation

After performing the heartbeat counting task, the thermocouples and additional electrodes for measuring electrodermal activity were put on. Participants received verbal instructions about the visuotactile stimulation procedure and were helped to put on the HMD. After reading instructions on the HMD, they performed a test trial where they selected “strongly agree” on a visual analogue scale (VAS) from “strongly disagree” to “strongly agree” to indicate that they were ready. A few seconds of exposure to a synchronous image of their own hands on the table followed to acquaint participants with the task and the virtual environment. For the experiment participants were instructed to not move and keep especially the head in a fixed position.

First a block with synchronous and asynchronous visuotactile stimulation of 60 s each was presented. Asynchrony was achieved by adding a 594 ms delay to the 139 ms intrinsic delay. The order of synchrony was counterbalanced across participants. After the 60 s of stimulation, the experimenter threatened the participant’s left hand with a plastic knife in a stabbing motion, which was followed by a 30 s rest period where the video feedback was displayed without any tactile stimulation, to assess change in heartrate. Participants were informed about the knife threat before starting the experiment, but did not know when it would occur. Both the synchronous and asynchronous condition were followed by the (dis)embodiment questionnaire. A block of 180 s of synchronous and synchronous visuotactile stimulation followed. After 180 s of visuotactile stimulation, the experimenter threatened the participant’s left hand with the plastic knife in a sliding motion. The 30 s rest period followed again. The 180 s blocks were aimed at assessing HRV during the manipulation of embodiment, and were not followed by the embodiment questionnaire. Again, the order of synchrony was counterbalanced across participants.

The experiment was concluded with a brief semi-structured interview on the experiences of the participant and a short debriefing. The full procedure took about 45 minutes.

#### 2.1.4 Measures of illusion strength

##### 2.1.4.1 (Dis)embodiment questionnaire

The subjective experience of the illusion was assessed with a questionnaire (see Table 1 for illusion related questions, an additional control question (q2, It seemed as if the seen hand resembled my own hand in terms of its shape and structure), and manipulation check (q3, It felt as if the stroking I felt on my hand was due to the seen stroking) were used), which was based on other studies, including the original rubber hand illusion (Botvinick & Cohen, 1998), the full-body illusion (Lenggenhager et al., 2007), the psychometric approach developed by Longo and colleagues (2008a), and additional new items to specifically assess disembodiment. Participants indicated on a VAS scale ranging from “completely disagree” to “completely agree” how much they agreed with each of the 11 statements. Based on the PCA (see section 2.2.1), three subscales of disownership, deafference, and embodiment were identified. The questionnaire was displayed in the HMD and participants responded by means of head movements.

**Table 1.** Factor loadings from the PCA on 9 items of the questionnaire in the asynchronous visuotactile condition.

|  |  | Varimax rotated factor loadings |  |  | commonalities |
| --- | --- | --- | --- | --- | --- |
|  | Sometimes I felt... | Component 1<br>*Disownership* | Component 2<br>*Deafference* | Component 3<br>*Embodiment* |  |
| q4 | alienation from my body | **.81** | .21 | .05 | .70 |
| q6 | as if my body does not belong to me anymore | **.78** | .18 | .29 | .73 |
| q9 | the seen body as an image rather than as my actual body | **.65** | .24 | -.04 | .50 |
| q10 | as if my body was numb | .12 | **.81** | -.03 | .68 |
| q8 | as though the experience of my hand was less vivid than normal | .23 | **.81** | -.14 | .73 |
| q7 | as though my body had disappeared | .25 | **.78** | .30 | .75 |
| q1 | as if the body I saw was my own | .02 | .00 | **.89** | .79 |
| q11 | as if I could move the seen body | .10 | .05 | **.81** | .67 |
| q5 | as if I was looking at another person's body | **.53** | -.05 | **.57** | .61 |
| Eigenvalues |  | 2.11 | 2.06 | 1.98 |  |
| % of variance |  | 23 | 23 | 22 |  |

*Note.* Factor loadings > .50 are in boldface.

##### 2.1.4.2 Skin temperature

Skin temperature was measured with an HH309A Data Logger thermometer (Omega, Stanford, CT, USA) at a 0.5 Hz sampling rate. Two thermocouples were placed on the left and right ventral side of the wrist, and a third on the back of the neck. A fourth thermocouple was used to monitor room temperature. Temperature was measured for the full length of the visuotactile stimulation in each condition. For each thermocouple, a baseline was calculated as the average temperature of the first 6 s of recording. This average value was subtracted from the subsequent recordings to represent the relative change in skin temperature across the stimulation period. In the short conditions, average temperature change was computed over 54 s (60 – 6 s baseline). In the long conditions, the average temperature change was computed over 174 s. One participant had to be excluded from the analyses due to technical problems. Two additional participants in the 60 s-blocks, and four in the 180 s-blocks were excluded due to missing data. We controlled for changes in room temperature by assessing differences in room temperature change between the asynchronous and synchronous condition, which were not significant in both the short block (*p* = .54) and the long block (*p* = .33).

##### 2.1.4.3 Skin conductance responses

Threat evoked SCRs were recorded with a Biopac MP150 system and EDA100C amplifier (Goleta, USA) at a 1000 Hz sampling rate. Two electrodes with electrode paste were placed on the participant’s index and middle finger of the non-stimulated right hand. The experimenters threatened the left hand of the participant, by making a stabbing motion in the short block, and a sliding motion in the long block, without touching the hand. A sound signal on the experimenter’s headphones indicated the onset of the threat, and a manual marker was placed in the raw data file immediately after presenting the threat. The data was processed in Acqknowledge software (Version 4.1, Biopac, Goleta, USA). The SCR was identified as the maximum amplitude in electrodermal activity around the threat event. It was expressed as a proportion of the average SCR, based on all four threat responses. Absent responses were registered as missing values. Data from five participants were excluded from the analysis due to missing responses or technical difficulties.

##### 2.1.4.4 Heart Rate Variability

A synchronous and asynchronous block of three-minute-long visuotactile stimulation was added to the procedure to assess HRV. ECG was recorded with the Biopac MP150 system as described previously. 160 seconds of recording were used, with an onset 10 s after the stimulation onset up to 10 s before the threat marker. The R-package RHRV (Rodriguez-Linares et al., 2017) was used to detect R-peaks and extract the Root Mean Square of the Successive Differences (RMSSD) as a measure of HRV. Data from four participants were excluded from the analysis due to technical difficulties.

##### 2.1.4.5 Data analysis

Data were analyzed with R (R Core Team, 2018) version 3.5.1. Alpha level was set at 0.05, or 95% confidence intervals, excluding 0, and p-values were adjusted for multiple comparisons using false discovery rate (FDR) corrections (Benjamini & Hochberg, 1995). Data were tested for normality, and appropriate tests were used accordingly. Details of preprocessing of the physiological data are described above.

##### 2.1.4.6 Principal Component Analysis

A PCA was used to investigate the structure of participants’ experience, and to quantify the complex experience during this illusion. The PCA was conducted on the questionnaire data after synchronous or asynchronous visuotactile stimulation. In order to maximize the number of participants we took the questionnaire data from Experiment 1 and questionnaire data from the long visuotactile stroking of Experiment 2 (see below) as well as additional data of 16 participants in an unpublished experiment (see supplementary material, Table S1, for descriptive statistics and item comparisons of these additional participants). Exposure time was 60 s in experiment 1 (*n* = 30), 90 s in experiment 2 (*n* = 32), and differed for the additional data between 60 s (*n* = 6) and 90 s (*n* = 9).

Two PCAs were separately run for the asynchronous and synchronous conditions. Adequacy of using PCA was assessed by Bartlett’s test of sphericity, which was highly significant for both the asynchronous (X^2^(55) = 238.6, *p* < .0001), and synchronous condition (X^2^(55) = 506.9, *p* < .0001), indicating that correlations between individual items were sufficiently large for PCA. The overall Kayser-Meyer-Olkin (KMO) measure verified that the sample size was adequate, both for the asynchronous (KMO = 0.71), and synchronous (KMO = 0.85) condition. The items for manipulation check (q2) and control item (q3) were excluded from the PCA, based on the low expected correlation with any of the other questionnaire items in the asynchronous conditions, as well as their poor individual KMO (both < 0.55) (Kaiser, 1974). An initial PCA was computed with 9 components. Inspection of the eigenvalues of each component and the scree plot (supplementary material, Figure S1) justified retaining three components for the secondary PCA.

### 2.2 Results

#### 2.2.1 Principal component analysis of questionnaire

The secondary PCA revealed three components, that together explained 68% of the variance in the questionnaire data (see Table 1 for component loadings after varimax rotation, and explained variance of each component). The first component we termed disownership and comprised items that refer to the experience of not belonging of the body, alienation, and perceiving the body as an image rather than an actual body (q4, q6, and q9). The second component was termed deafference (Longo et al., 2008) and included items related to the feeling of numbness, vividness, and disappearing of the body (q10, q8, q7). The final component, embodiment, consisted of items related to the experience of own body ownership, agency and looking at one’s own body (q1, q11, q5).

Responses to questionnaire items are in line with our hypotheses (Figure 2, see supplementary material Table S2 for descriptive statistics and results of the comparisons for all individual items). Participants report increased disownership, increased deafference and reduced embodiment after asynchronous visuotactile stimulation compared to synchronous stimulation. Responses to the control item (q3) did not differ between conditions, and the manipulation check item (q2) differed between the synchronous and asynchronous condition, which confirmed that participants were able to perceive the manipulation.

**Figure 2:**
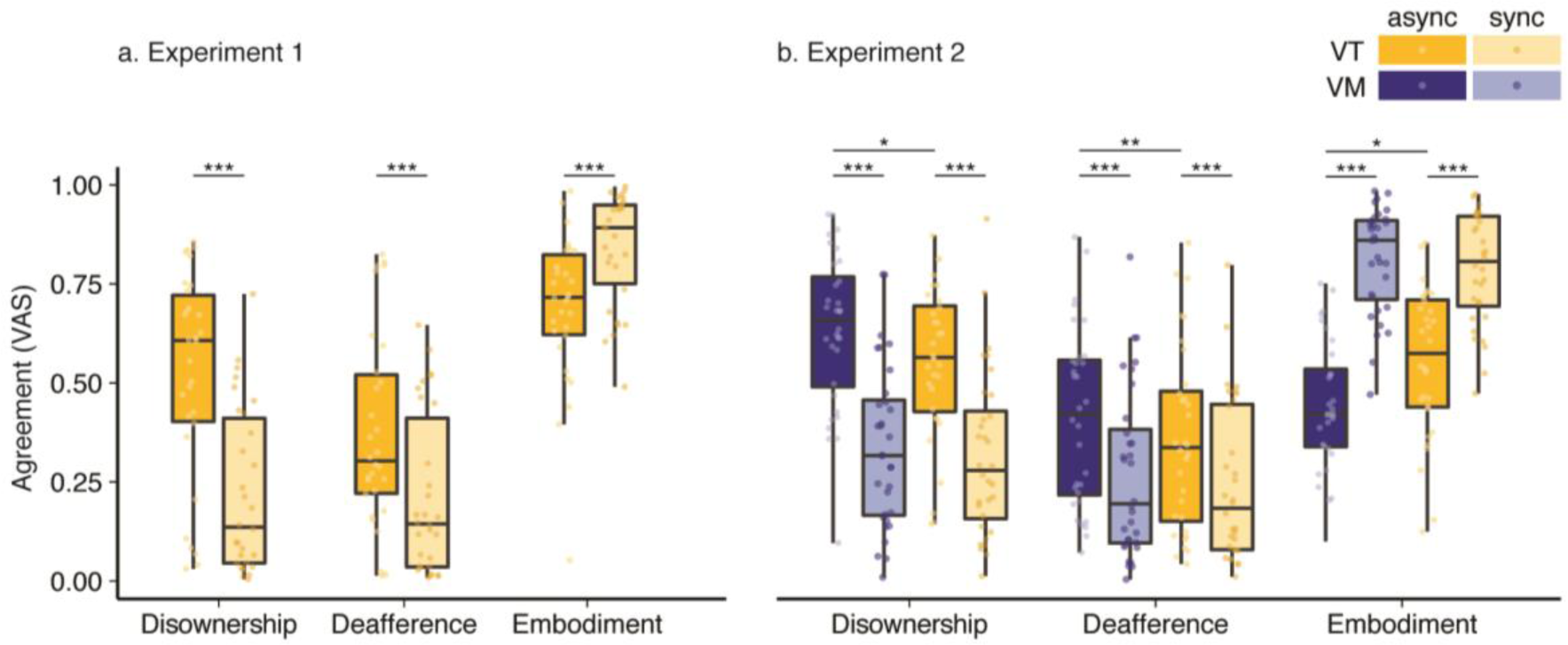
Questionnaire data, medians and interquartile ranges are displayed. The three components of the questionnaire differed significantly between the synchronous (sync) and asynchronous (async) visuotactile (VT) stimulation in Experiment 1 (a). In Experiment 2 (b) there were significant differences between the synchronous and asynchronous stimulation for both the visuotactile, and visuomotor (VM) stimulation, as well as between visuotactile and visuomotor stimulation in the asynchronous, but not the synchronous condition.

#### 2.2.2 Skin conductance responses

Previous studies demonstrated that SCRs to threats increased after synchronous stroking in rubber hand illusion like paradigms (e.g. Armel & Ramachandran, 2003; Petkova & Ehrsson, 2008), and one study showed that reduced SCR to a threat after multisensory mismatching stimulation (Gentile et al., 2013). As disownership was higher in the asynchronous condition, we hypothesized that SCRs would be reduced as compared to the synchronous condition. Even though we found an increase in skin conductance after threat, we did not observe significant differences between the synchronous and asynchronous condition in neither the short, (synchronous: *Mdn* = 1.13, IQR = 0.90 − 1.38; asynchronous: *Mdn* = 0.94, IQR = 0.71 – 1.16; *Z* = −1.42, *p* = .16) nor the long block (synchronous: *Mdn* = 1.01, IQR = 0.60 – 1.19; asynchronous *Mdn* = 0.94, IQR = 0.58 – 1.10; *Z* = −0.48, *p* = .63). This indicates that there was no difference in stress response to a threatening stimulus to the hand after asynchronous as compared to synchronous stimulation, even though participants subjectively experienced less embodiment, and increased disownership over their own hand.

#### 2.2.3 Skin Temperature

Comparisons between synchronous and asynchronous conditions in the short block did not reveal any significant differences between conditions in temperature change for the neck (*p* = .76), right hand (*p* = .38), or left hand (*p* = .27). However, in the long block, there was a significantly smaller increase in skin temperature of the left hand across the trial in the asynchronous (*Mdn* = 0.038, IQR = −0.009 – 0.143) than synchronous condition (*Mdn* = 0.078, IQR = 0.012 – 0.158; *Z* = −2.09, *p* = .04, *r* = −.30). There were no differences for the neck (*p* = .37) or the right hand (*p* = .72). We further aimed to disentangle this small, but significant effect for the left hand, by assessing differences between the conditions in each of the three minutes separately, but these analyses did not show any significant differences (all *p*s > .48).

#### 2.2.4 Heartrate variability

HRV, as quantified by the RMSSD, did not differ between the synchronous (*Mdn* = 30.62, IQR = 21.22 – 54.37) and the asynchronous condition (*Mdn* = 32.84, IQR = 24.61 – 45.30; *Z* = −0.25).

#### 2.2.5 Relation of illusion strength and interoceptive accuracy

Overall mean interoceptive accuracy was 0.62 ± 0.17, which is comparable to other studies (e.g. 4/2/2019 11:24:00 AM. We performed a median split on interoceptive accuracy scores to assess the differences in previously reported significant effects of synchrony between participants with high (*Mdn* = 0.77, IQR = 0.67 – 0.85) and low (*Mdn* = 0.46, IQR = 0.43 – 0.52) interoceptive accuracy. There was no significant difference between participants with high and low accuracy in the subjective strength of the illusion (difference between category average in synchronous and asynchronous) for the disownership (*Z* = −0.48, *p* = .63), deafference (*Z* = −0.33, *p* = .74), and embodiment (Z = −0.63, p = .53) category.

### 2.3 Discussion Experiment 1

In this first experiment we showed that asynchronously shown stroking of one’s own real hand using a video-based virtual reality setup leads as predicted to consistent and significant changes in the subjective sense of the bodily self as indexed by the response to the questionnaire. According to the principal component analysis the response to this questionnaire can be clustered in three main components, namely disownership, deafference and embodiment. During asynchronous as compared to synchronous stroking embodiment for one’s own body is reduced while the sense of disowernship and deafference are enhanced. In contrast to our prediction based on rubber hand illusion like setups, these changes were not reflected in the physiological response. The electrodermal response to threat remained equally high during both conditions, and the temperature measure, only showed a mild trend towards a lesser increase in temperature in the asynchronous condition. Furthermore, we did not find the predicted relation between the individual strength of interoception and the subjective measures of the illusion. Both these effects we relate to important conceptual and phenomenological differences between losing ownership for the real hand and extending ownership to an external object/body part (see section 4.3 in the main discussion).

## 3. Experiment 2

### 3.1 **Methods**

#### 3.1.1 Participants

Thirty-two healthy volunteers participated in Experiment 2 (7 males; 21.2 ± 3.9 years old). None of the participants took part in Experiment 1, and all gave informed consent and received either course credits or a financial compensation. The protocol was approved by the Ethics Committee of the Faculty of Arts and Social Sciences at the University of Zurich (Approval Number 17.12.15). The study was performed in accordance with the ethical standards of the Declaration of Helsinki.

#### 3.1.2 Apparatus for stimulation

The apparatus to present visual stimulation was identical to Experiment 1 (see 2.1.2). An additional laptop was used to play a metronome sound with its built-in speakers.

#### 3.1.3 Procedure

The experiment consisted of two different parts: first, two blocks with multiple trials of short stimulation, either visuotactile or visuomotor were presented, then four conditions of longer stimulations, either visuomotor or visuotactile both either synchronous or delayed were presented (see Figure 1b for an overview of the procedure). When participants were ready, they were helped to put on the HMD and read instructions on the screen. Similar to Experiment 1, the testing procedure was preceded by a test trial to practice giving responses on the VAS scale, and exposure to a synchronous image of the participant’s hands on the table for a few seconds.

For the visuotactile block, participants were asked to fix their left hand between the two markers on the table while they were stroked with a small paintbrush on their index and middle fingers. For the visuomotor block they were asked to move their left hand from the left to the right marker and back repeatedly, following the rhythm of a metronome (set to 1Hz). Each trial lasted 7 s and was followed by the question “Was the touch/movement you saw and felt synchronous?”, which could be answered by either selecting *yes* or *no*. This question was followed by the statement “It felt as if the hand that I saw was my own”, which could be answered on a VAS scale ranging from *strongly disagree* to *strongly agree.* The first two blocks consisted of 40 trials with four repetitions of 10 possible delay steps of 66 ms each, resulting in a range from 0 to 594 ms (plus the intrinsic 139.1 ms delay). The order of the visuomotor and visuotactile block were counterbalanced across participants.

Finally, a block of longer stimulation followed, where we presented four conditions (synchronous visuotactile, synchronous visuomotor, asynchronous visuotactile and asynchronous visuomotor) in counterbalanced order. The asynchronous conditions had a delay of 594 ms (plus the intrinsic 139.1 ms delay). During the visuomotor conditions, participants moved their hands as in the previous block but for a longer period; similarly, for the visuotactile condition, participants were stroked on their hand with a paintbrush randomly for a period of 90 s. After each condition, they were asked to answer the (dis)embodiment questionnaire (see section 2.1.4.1).

Participants could take breaks and remove the HMD in between blocks. The experiment was concluded with a brief semi-structured interview on the experiences of the participant and a short debriefing. The overall procedure took about 50 minutes.

#### 3.1.4 Measures of illusion strength

The assessment for the short stimulation was based on simultaneity judgment methods used to measure temporal windows of multisensory integration (Engel & Dougherty, 1971; Hirsh & Fraisse, 1964; Hoover & Harris, 2012, 2016), and an embodiment question derived from several studies (Botvinick & Cohen, 1998; Dobricki & Rosa, 2013; Lenggenhager et al., 2007). After each block of 90 s, participants completed an identical questionnaire as in Experiment 1. Item 2 differed between conditions, and was “It felt as if the movement I felt was due to the seen movement” in the visuomotor condition, and “It felt as if the stroking I felt on my hand was due to the seen stroking” in the visuotactile condition.

#### 3.1.5 Data analysis

The same software and parameters for significance were used as in Experiment 1. The questionnaire was analyzed using Wilcoxon signed-rank tests to assess the effect of synchrony (visuotactile synchronous vs. visuotactile asynchronous and visuomotor synchronous vs. visuomotor asynchronous) and the effect of modality (visuotactile synchronous vs. visuomotor synchronous and visuotactile asynchronous vs. visuomotor asynchronous).

Sensitivity to delay was assessed by determining the Point of Subjective Equality (PSE) for each participant in the visuomotor and visuotactile condition separately. To this end, logistic psychometric functions were fitted to the forced choice synchrony judgements of each participant, using a binomial Generalized Linear Model (glm) with delay as a predictor. The estimated coefficients of the glm were used to calculate the PSE: −β_0_ / β_1_, where β_0_ corresponds to the intercept and β_1_ to the slope. Goodness of fit was assessed with the Hosmer-Lemeshow test, and data of one participant in the visuotactile condition was excluded due to bad fit of the glm. All other psychometric curves did not yield a significant test result, and corresponding PSEs were thus used for further analyses.

Generalized linear mixed models were fitted with the lme4 package in R (Bates, Mächler, Bolker, & Walker, 2015). A generalized linear mixed model was fitted to the VAS ownership ratings in the short block, across different delays, which ensured for adequate power while considering the repeated measures within individuals. The intraclass correlation demonstrated that observations within individuals were non-independent (ICC(1) = .27, *F*(31, 2528) = 31, *p* < .001), thus justifying the use of a mixed model. Visual inspection of diagnostic plots of the residuals showed that these were normally distributed. The model that included both a random intercept and slope for individuals, where VAS ratings were explained as a function of delay, fitted the data better than the model that included only the random intercept and no random slope (X^2^(2) = 470, p < .001). Therefore, we used the random intercept and slope model for further hypothesis testing.

### 3.2 Results

#### 3.2.1 Questionnaire

To assess the subjective experience of participants after 90 s of visuotactile or visuomotor stimulation, differences between responses to questionnaire items in the asynchronous and synchronous conditions were assessed (see Figure 3; and supplementary material Table S3 and S4 for descriptive statistics and results for each individual item). The results confirmed our hypothesis that asynchronous visuotactile and visuomotor stimulation induces a feeling of disownership for the real hand, and followed a same pattern as in Experiment 1. There was a significant main effect of condition for the three illusion related factors that were determined in the PCA (see section 2.2.1). Interestingly, the reduction of embodiment and increase in deafference and disownership were stronger in the asynchronous visuomotor than visuotactile condition. There were no significant differences between conditions for the control item (q3) and the differences in q2 confirmed that participants were able to perceive the synchrony manipulation.

**Figure 3:**
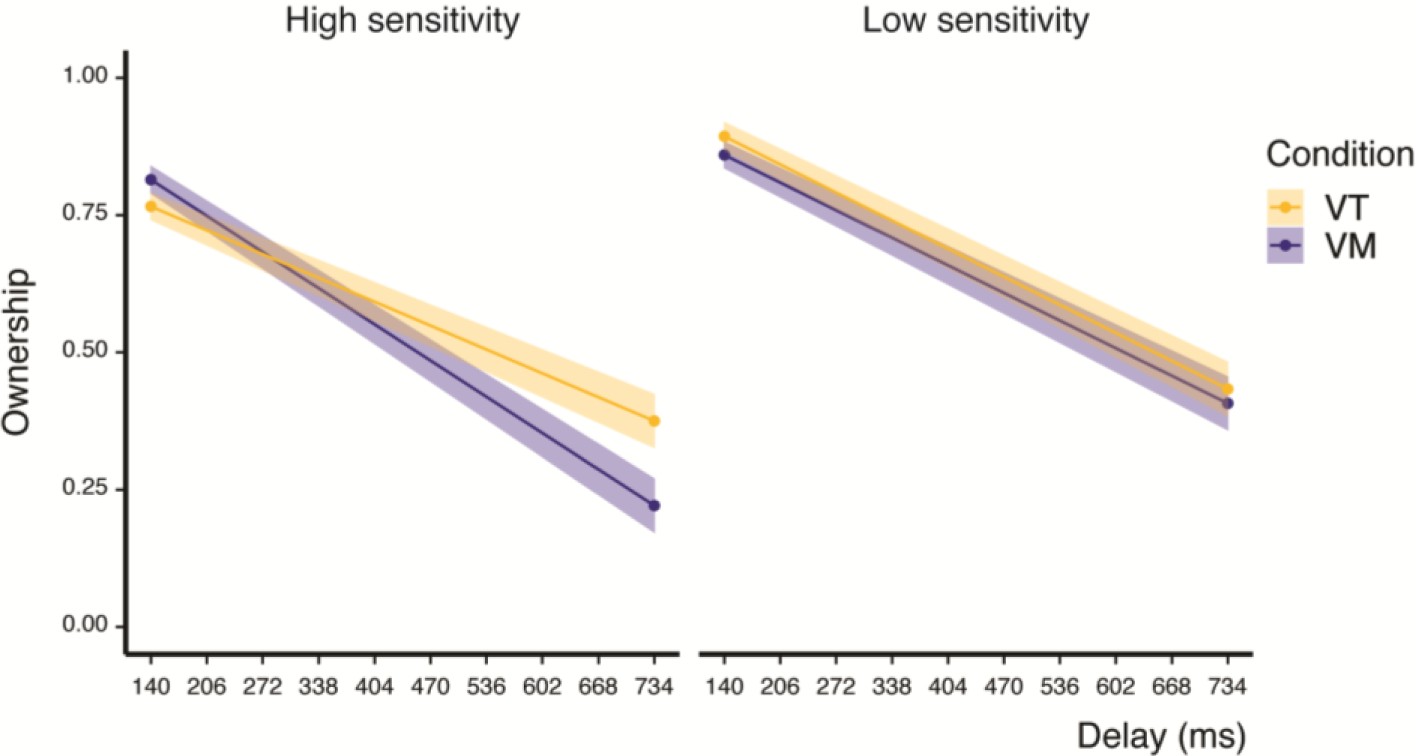
The three-way interaction of delay, modality (visuotactile (VT), and visuomotor (VM)) and sensitivity is displayed. Lines show predicted values from the model, where sensitivity was set to *M* − 1 *SD* for the high sensitivity group and *M* + 1 *SD* for the low sensitivity group.

#### 3.2.2 Synchrony judgements

To assess whether sensitivity to delay was affected by modality of stimulation, we compared the PSE in the visuomotor (*M* = 0.338, *SE* = 0.015) and visuotactile condition (*M* = 0.327, *SE* = 0.014). There was no significant difference between the two conditions (*Z =* −.55, *p* = .58). Sensitivity was also not correlated with relative changes in any of the questionnaire components between the synchronous and asynchronous stimulation in both the visuotactile (all *p*s > .59) or visuomotor (all *p*s > .44) condition.

#### 3.2.3 VAS illusion ratings after short stimulation

First, we fitted a model which included fixed effects for delay and condition, and their interaction (see supplementary material Table S5 for model coefficients). To explore whether sensitivity to delay for the different modalities, as quantified by the PSE, explained additional variance, we added the main effect and the two and three-way interactions with delay and condition in a second model. The model fit of the PSE-model was better than of the initial model (BIC model 1 = −1676.7, BIC PSE-model = −1709.5), and the PSE-model explained 32% of the variance in VAS ownership ratings (pseudo *R*^2^ = .32). Adding age as a predictor did not improve the model fit (BIC age-model: − 1705.2) and was thus removed from the model. There was a significant three-way interaction of all predictors (delay × modality × PSE; *b* = 2.19, 95% CI: 1.15, 3.23, *t* (2213.7) = 4.13, *p* < .001; see supplementary material Table S6 for all model coefficients). Overall, VAS ratings of ownership decreased with increasing delay. A stronger decrease in ownership was present especially in the visuomotor condition for participants with high sensitivity for delay. For lower sensitivity to delay there was no strong difference in strength of decrease between the visuotactile and visuomotor condition (see Figure 3).

### 3.3 Discussion Experiment 2

The results from the long stimulation in Experiment 2 show that visuomotor asynchrony when actively moving the hand in the same setup as in Experiment 1 also induces a decrease in embodiment coupled with an increase in disownership and sense of deafference. These changes were significantly stronger during visuomotor than during visuotactile mismatch. In line with this, the results from the short time exposure to various delays show that while increasing delay attenuates embodiment in both modalities, in participants with high delay sensitivity were visuomotor delays affected embodiment already at smaller delays. Together these results might suggest a stronger contribution of visuomotor as compared to visuotactile synchrony in maintaining embodiment of the own hand or/and a heightened sensitivity to mismatch during active body movements as compare to passive touch. The general reduction of ownership and the stronger decay in the visuomotor condition might suggest that, while efferent signals form stronger predictions on the sensory signals, mismatches to those predictions may result in a more salient phenomenology of disembodiment (see section 4.2).

## 4. Discussion

In two separate experiments and a PCA for a larger sample, we set out to assess how mismatching multimodal signals about one’s own body alter the sense of embodiment in healthy participants. For this, the participant’s hand was passively stroked or actively moved while seen from a first-person perspective on an HMD in a realistic video-based environment. The visual signals were either delayed (asynchronous; experimental condition) or presented simultaneously (synchronous; control condition) compared to the bodily signals (i.e. tactile or motor related). We used a (dis)embodiment questionnaire as well as physiological measures that have previously reported to correlate with body ownership (Experiment 1), and a series of synchrony and embodiment judgements across different visuotactile and visuomotor delays (Experiment 2). The two studies revealed three main findings. First, both visuotactile and visuomotor mismatch led to increased disembodiment, which predominantly involved the feelings of disownership, deafference, and embodiment (PCA results Experiment 1 and Experiment 2). Second, visuomotor delay when actively moving the hand led to a stronger feeling of disembodiment than visuotactile delay during passive touch. In participants with high delay sensitivity this was also evidenced by a steeper decay of body ownership with increased delay for visuomotor signals (Exp 2). Third, implicit measures of body ownership such as SCR and skin temperature where not modulated by the illusion (Experiment 1).

### 4.1 Multimodal temporal mismatches from the own hand alter the bodily self

Subjective changes in embodiment were measured with a questionnaire given to the participants after a stroking period. In line with previous studies (Gentile et al., 2013; Kannape et al., 2019) asynchronous stimulation generally reduced the feeling of embodiment, suggesting that synchronous multisensory inputs are not only crucial to induce embodiment over a supernumerary body (e.g. Botvinick & Cohen, 1998) but also to maintain the sense of embodying one’s own body. In a PCA based on the asynchronous visuotactile stroking, we identified three main factors of the subjective disembodiment experience. These are disownership corresponding to the experience of not belonging of the body, alienation, and perceiving the body as an image rather than as an actual body; deafference, which, in accordance to Longo et al. (2008a) includes numbness and vividness, plus in our case disappearance of the own body; and embodiment, consisting of the experience of body ownership, agency and the feeling of looking at one’s own hand. Our results show that both visuotactile and visuomotor mismatches lead to increased disownership, deafference and decreased embodiment respectively, when compared to synchronous stimulation.

In the case of synchronous stimulation, only two main factors were identified in the PCA, namely embodiment and disownership, together accounting for 71% of the variance (see supplementary material, Table S7). These results exclude the deafference component found for asynchronous stimulation. While this might be expected since our bodily experience is not generally accompanied by a sense of deafference, it should be noted that asynchronous signals not only led to a disruption of the components found for synchronous signals but also to a new phenomenological component (see Longo et al., 2008 for similar results using a rubber hand illusion). This suggests that disembodiment does not only vary along the dimensions of embodiment and disownership, but also includes a sense of deafference.

As mentioned in the introduction, the direct study of disembodiment in contrast to supernumerary embodiment is not trivial, as important conceptual (e.g. de Vignemont, 2011; Folegatti et al., 2009) and neuroanatomical (Martinaud et al., 2017; Zeller, Gross, Bartsch, Johansen-Berg, & Classen, 2011) differences between these two mechanisms have been suggested. Furthermore, there is only indirect, sparse, and non-conclusive evidence of supernumerary embodiment altering disembodiment (de Vignemont, 2011; Folegatti et al., 2009). Thus the currently most common way to study disembodiment, namely in RHI-like paradigms (Barnsley et al., 2011; Longo et al., 2008; Moseley et al., 2008) is problematic as it a) does not necessarily apply to some disturbances in body ownership, and b) may not actually induce the phenomena of interest. Our findings, on the contrary, demonstrate that it is possible to directly induce disembodiment at a phenomenological level by manipulating own-body related signals, potentially overcoming the abovementioned problems. Thus, the experimental manipulation proposed here, may be more transferable and ecologically valid for the study of disembodiment frequently manifested in psychological, psychiatric and neurological conditions (Aglioti et al., 1996; Brugger & Lenggenhager, 2014; Davidson, 1966; Sierra et al., 2005).

### 4.2 The effect of visuomotor as compared to visuotactile mismatch on the phenomenology of disembodiment

Our questionnaire data from Experiment 2 replicated and extended the findings of Experiment 1 by showing that both asynchronous visuotactile signals as well as asynchronous visuomotor signals lead to increased disembodiment. Moreover, prolonged asynchronous visuomotor signals had a stronger effect on disembodiment compared to that of visuotactile signals. While previous studies using foreign bodies or body parts have suggested that the tolerance for asynchronous visuotactile versus visuomotor stimulation during embodiment might differ (Kalckert & Ehrsson, 2012; Kokkinara & Slater, 2014; Roel Lesur et al., 2018; Tsakiris et al., 2006) and the specific contribution of actively moving on the bodily self has been intensively discussed (Grechuta et al., 2019; Pia et al., 2019) to our best of knowledge, this is the first time that this comparison is made by directly manipulating signals explicitly related to the own body. While it is known that in the clinical population both alterations in the sensory and motor systems might correlate with feelings of disembodiment, our results suggest that there may be a stronger contribution of the latter to disembodiment. On these lines, for example the rubber hand illusion has been related to activity in the premotor cortex (Ehrsson, Holmes, & Passingham, 2005; Ehrsson, Spence, & Passingham, 2004); and in clinical cases Burin et al. (2015) found that participants with left upper-limb hemiplegia experienced a greater rubber hand illusion in their affected hand when compared to both their unaffected hand and a control group, arguing that the reduction of efferent signals in these participants contributed to weakening their own body ownership, resulting in a more plastic sense of body. Our results further extend these findings showing that in healthy participants, breaking visuomotor synchrony facilitates the sense of disembodiment.

The data from the short trials of different delay steps might provide a more sensitive measure of the relation between small multimodal mismatches and its subjective interpretation and disembodiment. As hypothesized, the results generally showed better asynchrony detection and a decreased sense of ownership over one’s own body with increased delay. This finding was true for both the tactile and the motor modality and there were no significant differences in terms of perceived delay between multimodal couplings. This is surprising as previous literature suggested a greater delay sensitivity depending on the strength of efferent signals (Hoover & Harris, 2012; Lau, Rogers, Haggard, & Passingham, 2004; Winter, Harrar, Gozdzik, & Harris, 2008, the latter however without a statistically significant difference). A possible reason for this difference to previous literature is that our protocol might not have had a high enough temporal resolution to assess small differences in synchrony judgement, as previous literature has found it varying between 22 ms (Hoover & Harris, 2012) and 29ms (Winter et al., 2008). Moreover, theoretical models would suggest that with the presence of efferent signals, there would be a stronger expectation of afferent signals (Wolpert, 1997), thus affecting the perception of the afferent stimuli.

High sensitivity to delay, however, predicted overall lower ownership ratings, and especially in the visuomotor condition a faster decay. While previous studies have shown that greater temporal binding windows (TBW) of multisensory integration increase susceptibility to illusory embodiment of a rubber hand (Costantini et al., 2016) our results show that participants with high delay sensitivity have an overall stronger tendency to lose body ownership with increased delay between visuotactile or visuomotor signals than participants with lower sensitivity. A recent study found that the binding of incongruent multisensory signals in the ventriloquist effect (an effect where the location of an auditory stimulus is mapped to that of a visual stimulus: Pick, Warren, & Hay, 1969; Talsma, Senkowski, Soto-Faraco, & Woldorff, 2010) drops after active movements (Zierul et al., 2019); this is, incongruent signals are more easily bound when no efferent signals are involved. Zierul et al. (2019) expected, following a predictive coding account, that action would modulate the predictions and therefore bind incongruent stimuli more with action than without; however, their results showed the contrary. The authors thus hypothesize that action did form a *stronger* prediction, yet resulting mismatches were more salient and therefore multisensory incongruences were more evident. In our results, a similar explanation could be applied, i.e. expectations based on the motor-prediction were broken; while for the only visuotactile signals these expectations were not present. Moreover, in the visuomotor task, there is, next to matching between the motor command and the seen visual consequence, an additional mismatch of proprioceptive and visual signals that is not present during purely visuotactile tasks. This might explain a greater sensitivity to visuomotor delay as well as the steeper decay of body ownership. In this sense, the matching of motor predictions with their sensory consequences is not only important for the sense of agency, but seems to play an important role in the maintenance of a healthy sense of ownership (perhaps even more than the temporal coherence of somatosensory signals).

Importantly, low sensitivity to delay did not differently influence ownership sensation in the visuomotor and visuotactile tasks but rather generally predicted higher ownership. This could suggest that the effect might be mediated by stronger visual dependence: participants with stronger visual dependence would not be so sensitive to incongruencies to other senses since they rely stronger on vision as compared to other senses (Witkin & Asch, 1948). Indeed visual dependence has shown to be correlated with susceptibility to various multisensory illusions (David, Fiori, & Aglioti, 2014; Rothacher, Nguyen, Lenggenhager, Kunz, & Brugger, 2018). A stronger dependence on visual signals could thus explain why there was no difference in the decay of ownership for visuomotor and visuotactile tasks for participants with low delay sensitivity, however we did not objectively assess such dependence.

### 4.3 Physiological measures remain largely unchanged

The generally strong effect in the subjective measures of the illusion, was not mirrored in the here chosen implicit measures (skin temperature, SCR, HRV), where no or only rather weak effects were found. Only the temperature measure tentatively suggests a condition-specific effect by revealing a significantly smaller increase of temperature for asynchronous compared to synchronous stroking. This is in line with literature suggesting that a decrease in body temperature links to own-body disembodiment during illusory embodiment of a fake body (Moseley et al., 2008; Salomon, Lim, Pfeiffer, Gassert, & Blanke, 2013; but see also de Haan et al., 2017), or in neurological damage (Moseley et al., 2008; but see also Lenggenhager et al., 2015). As in previous literature, such relatively lower temperature was in our data specifically found for the stimulated hand (Macauda et al., 2015) and only after longer stimulation (cp. Macauda et al., 2015; Moseley et al., 2008, both reporting a drop in temperature only after more than a minute of stimulation), which might be related to the adaptation time homeostatic processes might need. However, when comparing temperature for different time periods of the long stimulation block, we found no significant differences between time periods. Thus, these results should be taken with caution. Moreover, an increasing amount of literature doubts a meaningful relationship between body ownership and body temperature (de Haan et al., 2017).

SCR is an indicator of physiological reactions to threat (Armel & Ramachandran, 2003; Ehrsson, 2007). Previous studies have linked embodiment of an external body part to a SCR when such body part is threatened (Armel & Ramachandran, 2003; Ehrsson, 2007), and one study has found a weaker SCR with decreased embodiment of the own body in a setup similar to ours (Gentile et al., 2013). Following this, we hypothesized to find a weaker response in the asynchronous compared to the synchronous stimulation condition. However, such an effect was not evident in our data, with both conditions showing a similarly strong response to threat. On the other hand, given that HRV has been suggested to be a measure of homeostatic processes (Berntson et al., 1997), we expected to find lower HRV during asynchronous stimulation due to a homeostatic disturbance, which was however not found.

So far, we can only speculate on the reasons for this lack of significant results in the chosen threat-related implicit measures. While generally the relationship between explicit and implicit measures of embodiment manipulations has been questioned (de Haan et al., 2017; Rohde, Luca, & Ernst, 2011; Rohde, Wold, Karnath, & Ernst, 2013), and in the case of HRV a recent study found no differences after altering embodiment in a full-body illusion (Park et al., 2016), it may be that the ecological congruency of the seen environment and body might have impeded an effect on our implicit measures. This is, in our setup participants are actually seeing their own hand and surroundings, with a higher degree of ecological plausibility compared to previous setups (e.g. Gentile et al., 2013). From an ecological point of view, it makes sense that participants would more readily extend the physiological reaction (protective space) to an external object than diminishing it. Furthermore, some of the described phenomenological alterations would not necessarily prevent a threat response, e.g. the feeling of deafference. Alternatively, it may be that even if there is disembodiment of one’s own body during asynchronous stimulation, it might be too fragile and that body perception may be immediately restored when attention is shifted away from the asynchronous stroking, regardless of limb-related multisensory synchrony. On these lines it has been proposed a low degree of ownership does not necessarily result in disownership, but that attention to the lack of ownership may (de Vignemont, 2011).

### 4.4 Interoceptive accuracy and its relation with bodily self plasticity

High interoceptive accuracy has previously been related to lower malleability of the bodily self in the context of the rubber hand illusion paradigm (Monti et al., 2019; Tsakiris et al., 2011). We thus expected interoceptive accuracy as measured by a heartbeat counting task to predict the degree of disembodiment after asynchronous stimulation. Yet, interoceptive accuracy did not predict the strength of disembodiment in the current study. Our findings are in line with recent studies showing no relation between interoception and suggestibility to bodily illusions (Crucianelli, Krahé, Jenkinson, & Fotopoulou, 2018; David et al., 2014). While future research is necessary to explain the difference between these studies, the fact that we did not evidence such correlation might again suggest differential mechanisms between extension of embodiment to an external object and reduction of own-body disembodiment. It thus raises the question whether there is a “general body plasticity” or if promptness to own-body disembodiment and to supernumerary embodiment may be separate components of such plasticity.

While such plasticity of the bodily self has traditionally been measured as the susceptibility to illusory supernumerary embodiment, there is currently no consensus on whether higher delay sensitivity in terms of own-body embodiment is a result of a more or less plastic bodily self or vice-versa. Costantini and colleagues (2016) found that a small TBW leads to lower susceptibility to illusory embodiment of a rubber hand, while we found that small TBWs lead to higher susceptibility to own-body disembodiment. This may seem paradoxical, since the same condition (small TBW) leads to both lower susceptibility to supernumerary embodiment and higher to own-body disembodiment. Such a contrast may suggest the need of separate components of bodily self plasticity, say one for supernumerary embodiment and one for own-body disembodiment. This would follow recent neuroanatomical findings in patients with disorders of embodiment (Martinaud et al., 2017; Zeller et al., 2011). This differentiation could help to explain the different results in implicit measures between previous literature and our study. However, it could also be that more proneness to a disembodiment illusion is actually a result of a less plastic bodily self, thus a weaker susceptibility to supernumerary embodiment. In this scenario, participants with a highly plastic bodily self would still maintain their sense of body even during stronger multimodal mismatches, adapting their bodily sense to the ongoing mismatching signals. While our data are inconclusive regarding this point, we propose that this is an important debate in the field of bodily self consciousness which in our view has not received enough attention. We hope to encourage future experimental inquiries that disentangle these questions, studies directly comparing our protocol with the rubber hand illusion may offer additional insights.

## 5. General Considerations, Challenges and Outlook

Our protocol offers methodological advantages for directly manipulating the perception of one’s own body, providing an easily replicable setup for studying and manipulating disembodiment without the need of involving supernumerary body (parts) and thus in a way that is more related to the loss of ownership described in certain psychiatric and neurological conditions. Moreover, the protocol used in Experiment 2 allows for a sensitive assessment of the contribution of various multimodal mismatches to the loss of body ownership and can be expanded to measure the temporal thresholds in relation to body ownership for other multimodal couplings. In contrast to illusory supernumerary ownership, which has described to occur after 11 s in visuotactile rubber hand setups (Ehrsson et al., 2004), 22.8 in active visuomotor (Kalckert & Ehrsson, 2017), and 36 in a visuotactile virtual hand setups (Perez-Marcos, Sanchez-Vives, & Slater, 2012) our Experiment 2 shows that even after short periods of stimulation (7 s), it is possible to manipulate the sense of one’s own body consistently and reliably (cp. also (Kannape et al., 2019). Such a procedure can be sensitive for comparing individual differences as well as between different populations.

Future studies comparing different populations and multisensory mismatch couplings are encouraged to shed light on the concept of bodily self plasticity. On these lines, a direct comparison of the rubber hand or a virtual hand illusion and our setup would offer important insights. It should be noted, however, that in our visuomotor task, participants were instructed to start and end every movement trajectory with their hand on the table during the visuomotor task, therefore the procedure also involved touch. Future studies should aim at constraining to the modalities in question.

With the increasing availability of mixed reality technologies, and in particular with the growing availability of augmented reality, understanding how our sense of body may change through our interactions with a mediated view of reality and the temporal mismatches that this may entail is of great importance. Again, the study of bodily self consciousness would benefit of studying more on how seeing one’s own body, instead of fake or virtual bodies, through digital visual manipulations affects embodiment. This is thus not only important at a theoretical and clinical level but may imply relevant practical knowledge for a near future where mixed reality technologies may be ubiquitous and thus constantly manipulate our sense of body.

## 6. Conclusion

The study of disembodiment is relevant for various clinical conditions and generally studied rather indirectly in the general population. Our disembodiment protocol may be important for the scientific study of bodily self consciousness, both to induce a sense of disembodiment and as an assessment tool. In particular, it may be a useful method to measure the degree and sensory weighting of bodily self plasticity in the general as well as clinical populations. The results of the two experiments presented here extend the previous literature showing that mismatching multisensory signals contribute to increased disembodiment of one’s own body as expressed by the phenomenological dimensions of disownership, deafference, and embodiment. Moreover they provide evidence for the differential contribution of sensorimotor signals compared to somatosensory in maintaining our sense of body. Lastly, we promote a debate regarding the concept of bodily self plasticity, proposing that either has two independent dimensions for supernumerary embodiment and for disembodiment respectively, or that strong susceptibility to disembodiment is a reflection of low bodily self plasticity.

## 7. Acknowledgements

M.R.L., M.L.W. and B.L. were supported by the Swiss National Science Foundation (grant number: 170511).

