## Supplementary material for "Achronopresence: how temporal visuotactile and visuomotor mismatches modulate embodiment"

**Supplementary figures and tables**

Table S1

*Descriptive statistics and pairwise comparisons of additional questionnaire data used in the PCA, N = 15.*

| | Visuotactile Synchronous | | Visuotactile Asynchronous | | | | |
|---|---|---|---|---|---|---|---|
| | Median | IQR | Median | IQR | *Z* | *p* | *r* |
| Disownership | 0.13 | 0.04 - 0.27 | 0.55 | 0.46 - 0.63 | -3.74 | <.001 | -.66 |
| q4 | 0.07 | 0.03 -0.16 | 0.67 | 0.38 - 0.76 | -4.01 | <.001 | -.71 |
| q6 | 0.10 | 0.03 -0.26 | 0.60 | 0.33 - 0.70 | -3.01 | .003 | -.53 |
| q9 | 0.17 | 0.03 - 0.39 | 0.58 | 0.37 - 0.70 | -2.71 | .007 | -.48 |
| Deafference | 0.04 | 0.03 - 0.11 | 0.58 | 0.27 - 0.63 | -3.52 | <.001 | -.62 |
| q10 | 0.06 | 0.02 - 0.13 | 0.72 | 0.36 - 0.77 | -3.33 | <.001 | -.59 |
| q8 | 0.06 | 0.03 - 0.10 | 0.53 | 0.09 - 0.67 | -2.64 | .008 | -.46 |
| q7 | 0.04 | 0.02 - 0.09 | 0.34 | 0.06 - 0.65 | -3.74 | <.001 | -.66 |
| Embodiment | 0.90 | 0.85 - 0.96 | 0.58 | 0.41 - 0.73 | -4.01 | <.001 | -.71 |
| q1 | 0.95 | 0.90 - 0.98 | 0.70 | 0.28 - 0.81 | -4.01 | <.001 | -.71 |
| q11 | 0.94 | 0.83 - 0.97 | 0.63 | 0.52 - 0.78 | -3.42 | <.001 | -.61 |
| q5 | 0.08 | 0.04 -0.29 | 0.61 | 0.31 - 0.83 | -3.33 | <.001 | -.59 |
| Control item and manipulation check | | | | | | | |
| q3 | 0.96 | 0.94 - 0.99 | 0.93 | 0.85 - 0.97 | -2.23 | .026 | -.39 |
| q2 | 0.96 | 0.84 - 0.98 | 0.68 | 0.24 - 0.72 | -2.57 | .010 | -.45 |

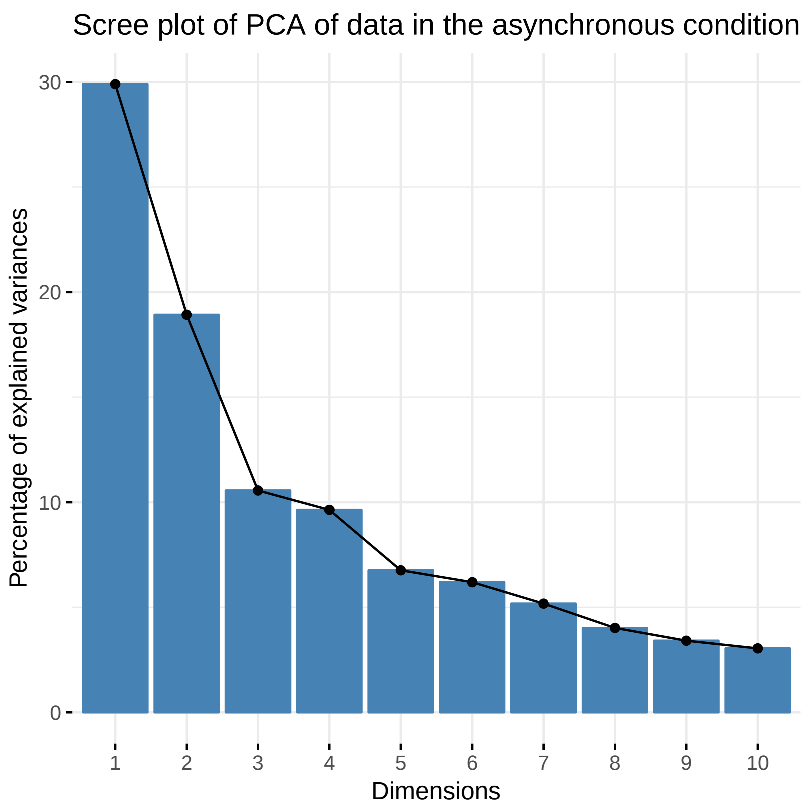


*Figure S1:* The scree plot of the PCA in the asynchronous condition justifies retaining three components for the secondary PCA.

Table S2

*Descriptive statistics and pairwise comparisons of questionnaire data in Experiment 1, N = 30.*

| | Visuotactile Synchronous | | Visuotactile Asynchronous | | | | |
|---|---|---|---|---|---|---|---|
| | Median | IQR | Median | IQR | *Z* | *p* | *r* |
| Disownership | 0.14 | 0.05 - 0.41 | 0.61 | 0.40 - 0.72 | -5.08 | <.001 | -0.66 |
| q4 | 0.11 | 0.03 – 0.30 | 0.59 | 0.25 – 0.78 | -4.59 | .005 | -0.59 |
| q6 | 0.07 | 0.03 – 0.23 | 0.54 | 0.18 – 0.70 | -4.96 | <.001 | -0.64 |
| q9 | 0.20 | 0.05 – 0.53 | 0.61 | 0.31 – 0.78 | -4.34 | <.001 | -0.56 |
| Deafference | 0.14 | 0.04 - 0.41 | 0.30 | 0.22 - 0.52 | -5.83 | <.001 | -0.75 |
| q10 | 0.11 | 0.03 – 0.43 | 0.29 | 0.12 – 0.69 | -4.17 | <.001 | -0.53 |
| q8 | 0.26 | 0.04 – 0.40 | 0.52 | 0.21 – 0.70 | -4.20 | <.001 | -0.54 |
| q7 | 0.07 | 0.03 – 0.17 | 0.18 | 0.07 – 0.46 | -4.34 | <.001 | -0.56 |
| Embodiment | 0.89 | 0.75 - 0.95 | 0.72 | 0.62 - 0.82 | -3.81 | <.001 | -0.49 |
| q1 | 0.91 | 0.88 – 0.98 | 0.78 | 0.67 – 0.86 | -4.17 | <.001 | -0.54 |
| q11 | 0.94 | 0.79 – 0.96 | 0.76 | 0.67 – 0.84 | -3.58 | <.001 | -0.46 |
| q5 | 0.10 | 0.04 – 0.28 | 0.27 | 0.13 – 0.62 | -2.78 | <.001 | -0.36 |
| Control item and manipulation check | | | | | | | |
| q3 | 0.93 | 0.84 – 0.97 | 0.91 | 0.81 – 0.97 | -1.31 | .191 | -0.16 |
| q2 | 0.85 | 0.57 – 0.97 | 0.51 | 0.28 – 0.75 | -3.28 | .001 | -0.42 |

Table S3

*Descriptive statistics of questionnaire data in Experiment 2, N = 32*

| | Visuotactile Synchronous | | Visuotactile Asynchronous | | Visuomotor Synchronous | | Visuomotor Asynchronous | |
|---|---|---|---|---|---|---|---|---|
| | Median | IQR | Median | IQR | Median | IQR | Median | IQR |
| Disownership | 0.28 | 0.16 - 0.43 | 0.57 | 0.43 – 0.70 | 0.32 | 0.17 – 0.46 | 0.66 | 0.49 – 0.77 |
| q4 | 0.17 | 0.09 - 0.36 | 0.6 | 0.34 - 0.69 | 0.2 | 0.10 - 0.30 | 0.72 | 0.56 - 0.82 |
| q6 | 0.21 | 0.09 - 0.29 | 0.62 | 0.32 - 0.73 | 0.18 | 0.07 - 0.33 | 0.69 | 0.57 - 0.85 |
| q9 | 0.5 | 0.24 - 0.68 | 0.67 | 0.40 - 0.76 | 0.58 | 0.26 - 0.74 | 0.62 | 0.33 - 0.80 |
| Deafference | 0.19 | 0.08 – 0.45 | 0.34 | 0.15 – 0.48 | 0.20 | 0.10 – 0.38 | 0.42 | 0.22 – 0.56 |
| q10 | 0.16 | 0.06 - 0.30 | 0.25 | 0.17 - 0.43 | 0.17 | 0.08 - 0.41 | 0.38 | 0.22 - 0.64 |
| q8 | 0.19 | 0.10 - 0.59 | 0.4 | 0.14 - 0.69 | 0.24 | 0.08 - 0.61 | 0.37 | 0.19 - 0.66 |
| q7 | 0.12 | 0.06 - 0.28 | 0.23 | 0.06 - 0.40 | 0.1 | 0.05 - 0.28 | 0.31 | 0.15 - 0.48 |
| Embodiment | 0.65 | 0.61 – 0.69 | 0.56 | 0.51 – 0.65 | 0.66 | 0.62 - 0.68 | 0.54 | 0.44 - 0.65 |
| q1 | 0.9 | 0.81 - 0.97 | 0.65 | 0.41 - 0.78 | 0.91 | 0.81 - 0.95 | 0.42 | 0.28 - 0.68 |
| q11 | 0.85 | 0.78 - 0.93 | 0.7 | 0.37 - 0.82 | 0.89 | 0.80 - 0.96 | 0.54 | 0.29 - 0.69 |
| q5 | 0.19 | 0.09 - 0.41 | 0.57 | 0.34 - 0.74 | 0.19 | 0.08 - 0.39 | 0.69 | 0.46 - 0.85 |
| Control item and manipulation check | | | | | | | | |
| q3 | 0.91 | 0.84 - 0.97 | 0.86 | 0.80 - 0.94 | 0.89 | 0.78 - 0.97 | 0.86 | 0.68 - 0.96 |
| q2 | 0.85 | 0.57 - 0.95 | 0.53 | 0.18 - 0.69 | 0.89 | 0.77 - 0.95 | 0.66 | 0.56 - 0.81 |

Table S4

*Results of Friedman tests and post-hoc comparisons of questionnaire in Experiment 2, using Wilcoxon Signed-Rank tests, FDR corrected p-values*

| | Friedman test | | | VTsyn - VTasyn | | | VMsyn - VMasyn | | | VTsyn - VMsyn | | | VTasyn - VMasyn | | |
|---|---|---|---|---|---|---|---|---|---|---|---|---|---|---|---|
| | $\chi^2$ | df | *p* | *Z* | $p_{corrected}$ | *r* | *Z* | $p_{corrected}$ | *r* | *Z* | $p_{corrected}$ | *r* | *Z* | $p_{corrected}$ | *r* |
| Disownership | 35.51 | 3 | <.001 | -4.42 | <.001 | -0.55 | -5.12 | <.001 | -0.64 | -0.05 | .963 | -0.01 | -2.31 | .021 | -0.29 |
| q4 | 39.38 | 3 | <.001 | -4.19 | <.001 | -0.52 | -5.09 | <.001 | -0.63 | -0.06 | .949 | -0.01 | -2.52 | .016 | -0.31 |
| q6 | 43.46 | 3 | <.001 | -4.72 | <.001 | -0.59 | -5.30 | <.001 | -0.66 | -0.25 | .803 | -0.03 | -3.04 | .003 | -0.38 |
| q9 | 7.76 | 3 | .051 | | | | | | | | | | | | |
| Deafference | 26.78 | 3 | <.001 | -3.36 | <.001 | -0.42 | -3.77 | <.001 | -0.47 | -0.69 | .488 | -0.09 | -1.98 | .048 | -0.25 |
| q10 | 28.56 | 3 | <.001 | -3.63 | .001 | -.045 | -3.41 | .001 | -0.43 | -1.71 | .087 | -0.21 | -3.50 | .001 | -0.44 |
| q8 | 5.66 | 3 | .129 | | | | | | | | | | | | |
| q7 | 19.95 | 3 | <.001 | -2.16 | .062 | -0.27 | -4.72 | <.001 | -0.59 | -0.10 | .919 | -0.01 | -1.94 | .069 | -0.24 |
| Embodiment | 59.36 | 3 | <.001 | -4.58 | <.001 | -0.57 | -5.92 | <.001 | -0.74 | -0.97 | .331 | -0.12 | -2.85 | .005 | -0.36 |
| q1 | 55.65 | 3 | <.001 | -4.19 | <.001 | -0.52 | -5.86 | <.001 | -0.73 | -0.05 | .963 | -0.01 | -2.54 | .015 | -0.05 |
| q11 | 52.46 | 3 | <.001 | -4.24 | <.001 | -0.53 | -6.23 | <.001 | -0.78 | -1.54 | .139 | -0.19 | -1.48 | .139 | -0.19 |
| q5 | 49.24 | 3 | <.001 | -3.63 | .001 | -0.45 | -4.90 | <.001 | -0.61 | -0.18 | .861 | -0.02 | -2.76 | .008 | -0.34 |
| Control item and manipulation check | | | | | | | | | | | | | | | |
| q2 | | | | -3.61 | <.001 | -0.45 | -3.68 | <.001 | -0.46 | | | | | | |
| q3 | 9.19 | 3 | .027 | -1.96 | .100 | -0.24 | -2.46 | .056 | -0.31 | -0.05 | .963 | -0.01 | -1.69 | .121 | -0.21 |

Table S5

*Summary of the initial mixed model, including the fixed effects for delay and condition, and their two-way interaction*

| | | Confidence interval | | | | | |
|---|---|---|---|---|---|---|---|
| fixed effects | *b* | lower | upper | *SE* | *df* | *t* | *p* |
| intercept | 0.923 | 0.866 | 0.981 | 0.029 | 46.29 | 31.49 | <.001 |
| Delay | -0.718 | -0.772 | -0.665 | 0.027 | 2525 | -8.84 | <.001 |
| Modality | 0.038 | 0.002 | 0.074 | 0.018 | 2525 | 2.29 | .04 |
| Delay x Modality | -0.154 | -0.230 | -0.079 | 0.039 | 2525 | -4.47 | <.001 |

*Notes: Modality (0 = Visuotactile, 1 = Visuomotor)*

Table S6

*Summary of the final mixed model, including the predictors delay, condition, and PSE, and the two-way and three-way interactions.*

| | | Confidence interval | | | | | |
|---|---|---|---|---|---|---|---|
| fixed effects | *b* | lower | upper | *SE* | *df* | *t* | *p* |
| intercept | 0.830 | 0.788 | 0.871 | 0.027 | 37.23 | 34.32 | <.001 |
| Delay | -0.717 | -0.880 | -0.553 | 0.082 | 33.82 | -8.69 | <.001 |
| Modality | 0.007 | -0.017 | 0.031 | 0.017 | 2473.88 | 1.84 | .066 |
| PSE | 0.790 | 0.485 | 1.099 | 0.210 | 1393.73 | 4.24 | <.001 |
| Delay x Modality | -0.164 | -0.232 | -0.096 | 0.035 | 2471.39 | -4.73 | <.001 |
| Delay x PSE | -0.724 | -1.609 | 0.149 | 0.446 | 1633.18 | -1.62 | .105 |
| Condition x PSE | -0.511 | -0.878 | -0.148 | 0.251 | 2086.97 | -3.26 | .001 |
| Delay x Modality x PSE | 2.188 | 1.153 | 3.233 | 0.530 | 2213.69 | 4.13 | <.001 |

*Notes: Modality (0 = Visuotactile, 1 = Visuomotor), PSE was mean-centered around 0, lower values correspond to high sensitivity, and higher values to low sensitivity.*

Table S7

*Results of the PCA on questionnaire responses in the synchronous condition*

| | Varimax rotated factor loadings | | |
|---|---|---|---|
| | Component 1 *Disownership* | Component 2 *Embodiment* | commonalities |
| q4 | **0.87** | 0.27 | 0.83 |
| q10 | **0.87** | 0.06 | 0.76 |
| q7 | **0.81** | 0.27 | 0.73 |
| q6 | **0.79** | **0.44** | 0.82 |
| q8 | **0.72** | 0.38 | 0.66 |
| q9 | **0.71** | 0.41 | 0.67 |
| q5 | **0.67** | 0.12 | 0.46 |
| q11 | 0.10 | **0.87** | 0.77 |
| q1 | 0.34 | **0.72** | 0.64 |
| Eigenvalues | 4.39 | 1.95 | |
| % of variance | 49 | 22 | |
